# Splicer: Phylogenetic Placement in Sub-Linear Time

**DOI:** 10.64898/2026.02.10.705130

**Authors:** Alexey Markin, Tavis Anderson

## Abstract

**Motivation:** Phylogenetic placement is an established approach for rapidly classifying new genetic sequences and updating a phylogeny without fully recomputing it. Popular maximum-likelihood placement methods, such as pplacer and EPA-ng, tend to struggle computationally when the size of the reference tree increases to tens or hundreds of thousands of sequences. As a more scalable alternative, distance-based and parsimony-based placement methods were introduced such as UShER. These methods, in principle, scale linearly as the size of the reference tree grows. However, as the scale of genetic and genomic sequences continues to grow nearly exponentially, developing algorithms that can perform placement in *sub-linear* time while maintaining accuracy becomes more crucial.

**Results:** Here, we develop *Splicer*, the first such algorithm that can perform placement in guaranteed 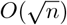 time. To achieve this performance, Splicer first decomposes the original reference tree into ‘blobs’ and constructs a phylogenetic scaffold tree linking representatives from different blobs. Every blob in such decomposition has at most 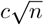 taxa, and the scaffold tree has at most 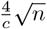 leaves, where *c* is any constant. Then, given the query sequences for placement, they are first placed onto a scaffold tree using pplacer or EPA-ng, and then placed more precisely within the respective blobs. We demonstrate the high accuracy of Splicer on an empirical influenza A virus dataset that has sparse coverage due to limited genomic surveillance. We also show that Splicer can, for the first time, apply maximum-likelihood placement to COVID-19 pandemic-scale data using a dataset with over 12 million SARS-CoV-2 reference genomes. Splicer scales the highly accurate maximum-likelihood approaches implemented in pplacer and EPA-ng to trees with millions of taxa and eliminates the necessity to curate and subsample genomic datasets for real-time classifications.

**Availability and implementation:** Splicer tool and source code are freely available at https://github.com/flu-crew/splicer.

## 1 Introduction

Phylogenetic placement is a common approach for updating a pre-built phylogeny and classifying new genetic or genomic sequences using a reference dataset [8, 1]. Phylogenetic placement tools attempt to find the most likely branch on the reference phylogeny where the new query sequence or sequences should appear if the full tree were to be newly inferred [15, 6]. Two of the most popular tools for phylogenetic placement are *pplacer* [14] and EPA-ng [5] which apply the maximum likelihood framework for identifying the placement branches for query sequences. Unfortunately, as the volume of data and the size of a reference phylogeny grow, these methods can become prohibitively slow on datasets with tens of thousands of reference genomes and may fail to accurately place queries. To overcome this bottleneck, novel placement methods that use genetic distance (e.g., APPLES [4]) or parsimony (UShER [20]) have been proposed. UShER was specifically designed to tackle the immense challenge of updating the SARS-CoV-2 phylogeny with millions of novel genomes being sampled annually at the peak of the COVID-19 pandemic. The continued use of UShER has demonstrated that it is well-suited for phylogenetic placement, and it is particularly strong when the sampling density of reference genomes is high.

Alternatively, several algorithms have been proposed to scale maximum likelihood phylogenetic placement to trees with hundreds of thousands of taxa. The most notable examples are pplacerDC [10] and SCAMPP [21]. The pplacerDC approach uses a divide-and-conquer method to place a new sequence on a backbone tree, whereas SCAMPP ‘carves out’ a subtree where the sequence can be placed based on sequence similarity. Both approaches have been successful in scaling maximum-likelihood placement to larger datasets. However, both of these approaches have an inherent limitation of a requirement to parse the full reference tree/dataset for each new placement run.

In this work, we propose a decomposition-based approach, *Splicer*, that can perform each placement in guaranteed *sub-linear* time. The key advantage of sub-linear algorithms is that they continue to scale well even as the size of reference datasets grow rapidly. In particular, Splicer guarantees that each placement will be performed in 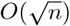 time and will be correct, assuming the underlying placement algorithm is accurate. Splicer achieves this by decomposing the reference dataset, i.e., a phylogenetic tree with an alignment, into connected subtrees called ‘blobs’ of size at most 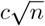, where *n* is the total number of reference sequences and *c* is some constant; e.g., *c* = 2. Then, each blob is collapsed into a single node on the reference phylogeny, and representative taxa are selected from some of the blobs, resulting in the *scaffold tree* (Figure 1A). The algorithm selects scaffold taxa in a way that ensures that every blob is represented by at least one node on the scaffold tree, maintaining the evolutionary history of the taxa. The decomposition algorithm runs in *O*(*n* log *n*) time and needs to be performed only once. After the decomposition is done, for each placement query, Splicer executes pplacer or EPA-ng on the scaffold tree to find approximate locations for the query sequences, and then places the query sequences more precisely within the respective blobs. At every step during placement, Splicer requires only 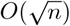 time. Additionally, we prove that if the underlying placing algorithm is accurate, then Splicer is guaranteed to be accurate as well (Theorem 2). To make Splicer applicable to sequence classification tasks, we implemented the capability to parse a clades/taxonomy definition file during the decomposition step, to then immediately classify query sequences during the placement step.

**Figure 1.**
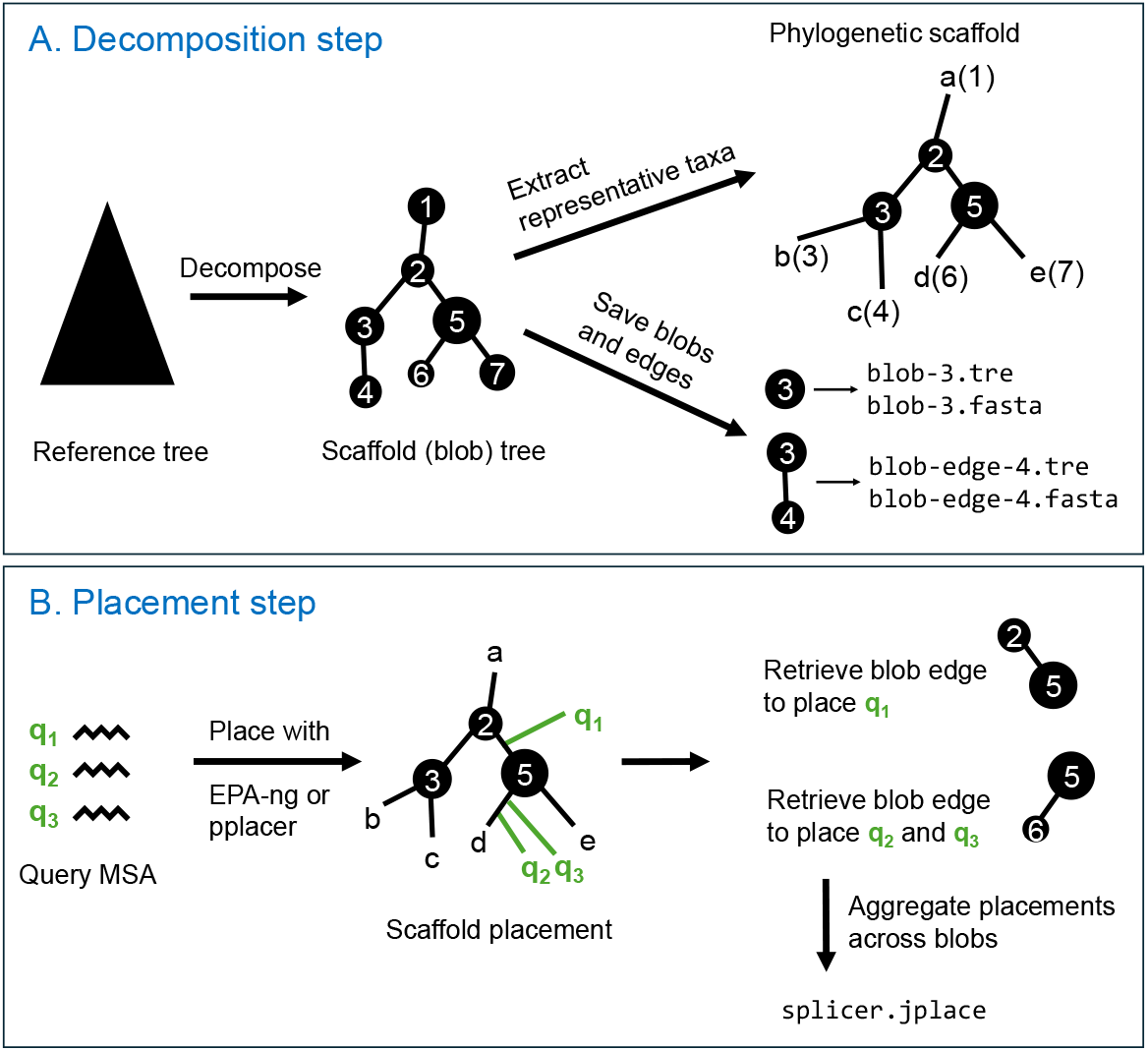
The overview of the decomposition algorithm of Splicer (A) and the sub-linear placement algorithm on the decomposed tree (B).

Using a large influenza A virus (IAV) dataset of 14,000 hemagglutinin (HA) sequences, we demonstrate that Splicer was the most accurate method, slightly outperforming UShER and SCAMPP. When placement was performed in bulk, Splicer+EPA-ng was the fastest method, spending only 0.2 seconds per sequence with this dataset. We compared the performance of the base methods (pplacer and EPA-ng) on a subset of 483 representative sequences selected from the 14,000 IAV sequences against the performance of other scalable methods (Splicer, UShER, and SCAMPP) using all available sequences. We demonstrated that utilizing the full reference dataset substantially enhances placement accuracy, reducing the error rate from 4% for EPA-ng to only 0.4% for Splicer+EPA-ng (Figure 4). This study suggests that using scalable placement methods on larger datasets is advantageous over subsampling strategies in terms of both speed and accuracy. Furthermore, to demonstrate the strong scalability of our method, we applied Splicer paired with EPA-ng to the SARS-CoV-2 genome dataset, which contains over 12 million reference sequences. Splicer performed bulk placement of 1000 genomes within 28 hours (100 seconds per sequence), scaling maximum-likelihood placement to pandemic-scale datasets for the first time.

## 2 Methods

### 2.1 Splicer algorithm

The Splicer algorithm consists of two steps. First, Splicer decomposes the reference tree and alignment with *n* taxa into ‘blobs’ of a guaranteed small size and constructs a scaffold tree representing the tree of blobs^1^. This step ensures that every subsequent placement query is processed in sublinear time, specifically 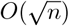. For given query sequences, Splicer first identifies blobs relevant to the queries by running pplacer or EPA-ng on the scaffold tree, and then it finds optimal placements within the identified blobs. Below, we describe the decomposition and placement algorithms in more detail and prove the sub-linear runtime of the placement. Finally, we demonstrate that if an underlying placement algorithm is consistently accurate, then Splicer is also guaranteed to be accurate.

#### 2.1.1 Reference tree decomposition

Let *T* be the reference phylogenetic tree with *n* leaves bijectively labeled by a taxon set *L*, and let ⟨*R*⟩ denote the multiple sequence alignment of the reference taxa.

We construct a scaffold tree *S*, where each node *v* in *S* represents a blob *B*(*v*) (a connected subtree of *T* ) such that

i. Every blob *B*(*v*) has at most 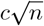 nodes, where *c >* 0 is some constant;
ii. Blobs *B*(*v*) are non-overlapping for different *v* ∈ *S*;
iii. Every node of *T* is represented by at least one of the blobs in *S*;
iv. Nodes *v*_1_ and *v*_2_ of *S* are connected if and only if there exists *w*_1_ ∈ *B*(*v*_1_) and *w*_2_ ∈ *B*(*v*_2_) that are connected in *T* .
v. Every node *v* of *S* has degree at most 3.

We construct *S* by iteratively merging adjacent nodes of *T*, and we make sure that no node in *S* has degree more than three at any point during the construction.

1. Initialize *S* to be the original tree *T* . That is, initially, each node of *S* represents the respective node of *T* . Let the *size* of a node *v* of *S* denote the number of leaves of *T* that it represents. We denote the size of *v* by |*v*|. Then, initially, each leaf node of *S* has size 1 and each internal node has size 0.
2. An edge (*u, v*) of *S*, where *v* is a child of *u*, can be contracted if (i) |*v*| *>* 0, (ii) 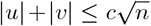, and (iii) the degree of *u* or *v* is less than 3.
3. Let *E*_*C*_ denote the set of all contractable edges of *S*. We maintain a priority queue of *E*_*C*_ ranked by the edge length (shorter edges are contracted first).
4. Contract a shortest edge (*u, v*) in *E*_*C*_. That is, we merge *u* and *v* into a new node *w* with |*w*| = |*u*| + |*v*|. Finally, update the priority queue by evaluating whether the edges incident on *u* and *v* can be contracted after the merge.
5. Repeat Step 4 until there are no contractable edges.

It is not difficult to see that these steps produce a scaffold tree *S* that satisfies properties (i)-(v) above. The runtime of this algorithm is *O*(*n* log *n*) because of the need to maintain a priority queue for edges that can be contracted. While it is possible to implement such a blobification algorithm without a priority queue and using only linear time, we believe that prioritizing shorter edges for contraction is important to make sure that the generated blobs represent meaningful clusters of sequences. Next, we prove that the number of blobs we get from the algorithm is guaranteed to be small.

##### Theorem 1.

*The scaffold tree S has at most* 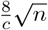 *nodes. Additionally, at most* 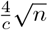 *of nodes have a degree less than 3*.

*Proof*. We classify all nodes of *S* into *small* and *large* nodes based on their blob size. A node *v* is small if 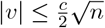, and otherwise it is large. Note that there are less than 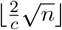 large nodes.

By *s*_1_, *s*_2_, *s*_3_, we denote the number of small nodes of degree 1 (leaves), degree 2, and degree 3, respectively. Similarly, by *l*_1_, *l*_2_, *l*_3_, we denote the number of large nodes of degree 1, 2, and 3, respectively. Further, *s* = *s*_1_ + *s*_2_ + *s*_3_ and *l* = *l*_1_ + *l*_2_ + *l*_3_ are the total numbers of small and large nodes in *S*. Note that a parent of a small node of degree 2 or 1 can only be a large node (otherwise, the two nodes can be merged into one). Then,

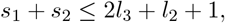

as degree-3 large nodes can parent at most two small nodes, and *l*_2_ nodes can parent at most one small node; plus, we need to account for the possibility of the root node being small. Then, the total number of degree-1 and degree-2 nodes is

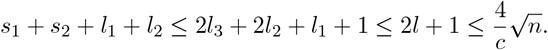

Next, we bound *s*_3_. Note that a child of an *s*_3_ node is either a large node or another degree-3 small node. That is, on a path from an *s*_3_ node to any leaf below it, there must be at least one large node. Then, consider the minimum subtree of *S* that contains all the large nodes, denoted *S*_*l*_. *S*_*l*_ must contain all degree-3 nodes (both small and large), and the leaves of *S*_*l*_ are only large nodes. The number of degree-3 nodes in such a tree is at most *l* − 1. Therefore, *s*_3_ ≤ *l* − 1, and

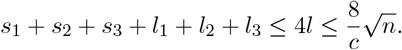

□

The next step is to transform the scaffold tree into a phylogenetic tree, so that query sequences can be placed onto it. To do that, we choose a single taxon closest to the root node from each of the degree-2 and degree-1 blobs (Figure 1A) to represent that blob. The subtree of *T, P*, induced by these taxa, then represents the phylogenetic scaffold tree. Every internal node of *P* corresponds to a unique blob. Additionally, the leaves drawn from degree-1 blobs represent these blobs uniquely as well. Note that the number of leaves in *P* is at most 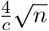 by Theorem 1.

The final step of the decomposition is writing the phylogenetic scaffold tree, the blob trees, and the *blob-edges* to disk. Note that blob-edges are two blobs connected by an edge merged into one tree. In theory, we want to choose the constant *c* so that the scaffold tree, individual blobs, and blob-edges are as small as possible. Note that the size of a blob-edge is upper bounded by 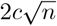. Then, choosing 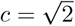 optimizes the balance between the scaffold tree size and the blob-edge size, making sure that we never need to process a tree of size larger than 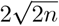. However, in practice, we found that *c* = 0.5 generally gives higher placement accuracy than 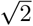.

##### Completing blobs

Each blob in the decomposition has at most three external edges connecting it to other blobs (up to two child links and one parent link). To make sure that we maintain all internal nodes of the blob when extracting it from the larger tree, we add a leaf to the blob for each of the external links.

In particular, for every internal node *v* ∈ *T* (i.e., excluding the leaves and the root), we define *c*^↑^(*v*) to be the closest leaf to *v* in *T* that is below *v*. Further, we define *c*^↓^(*v*) to be the closest leaf to *v* in *T* such that the path from *c*^↓^(*v*) to *v* contains at least one node above *v*. Note that *c*^↑^(*v*) and *c*^↓^(*v*) can be precomputed in linear time over all internal nodes of *T* using one postorder and one preorder traversals.

For a blob *B* and an edge (*b, e*) where *b* ∈ *B* and *e* ∉ *B* is external, we add leaf *c*^↑^(*e*) to the leafset of *B*. Similarly, for an external edge (*e, b*), we add the leaf *c*↓(*b*) to the leafset. For a blob-edge combining blobs *B*_1_ and *B*_2_, we perform a similar procedure, adding at most 4 external leaves for that pair.

##### Empty blobs

Some degree-3 blobs may contain no taxa from the reference tree - we refer to such blobs as *empty*. While rare in practice, here we define blob-edges when one of the blobs is empty. Assume that *B* is a blob with at least one taxon and let *B*_0_ be its neighbor blob that is empty.

If *B*_0_ is a parent of *B*, we define the blob-edge (*B*_0_, *B*) to be *B* with an additional edge leading to *c*^↓^(*ρ*), where *ρ* is the root of *B*. If *B*_0_ is a child of *B*, let *v* denote the node in *B* that is the parent of the root of *B*_0_. We define the blob-edge (*B, B*_0_) to be *B* with *c*^↑^(*v*) added as a child of *v*. This is done to make sure that all edges of the original tree are accounted for across the blob-edge trees.

If both *B* and *B*_0_ are empty, we do not create a blob-edge for that pair.

###### Lemma 1.

*Every edge of the original tree T is contained either in the scaffold tree, a blob, or a blob-edge*.

*Proof*. For an edge (*u, v*), if *u* and *v* belong to the same blob, then that edge will be represented by that blob by construction, as we make sure that all leaves and internal nodes of the blob are maintained. Assume now that *u* and *v* belong to different blobs *B*_1_ and *B*_2_. If *B*_1_ and *B*_2_ are both empty, then the edge (*u, v*) must be present in the scaffold tree. Otherwise, the edge (*u, v*) will be present in the edge-blob {*B*_1_, *B*_2_} by construction.

□

#### 2.1.2 Sub-linear placement algorithm

Let ⟨*Q*⟩ = (*q*_1_, …, *q*_*m*_) denote the query sequence alignment to be placed on the reference tree. Splicer first places the query sequences on the scaffold tree and then determines more precise placement edges by placing the query sequences within the relevant blobs or blob-edges (Figure 1B). Splicer uses pplacer or EPA-ng as the underlying placement algorithm at every step.

##### Placement on the scaffold tree

Splicer first places the query sequences onto the phylogenetic scaffold tree from the decomposition step. Recall that every edge in the scaffold tree either represents a pair of blobs (a blob-edge) or a single blob. That is, for a query *q* placed on a scaffold edge (*u, v*): if *B*(*u*) = *B*(*v*) then *q* gets assigned to the blob *B*(*u*), and otherwise it gest assigned to the blob-edge {*B*(*u*), *B*(*v*)}.

Splicer identifies the set of blob-edges *E* and the set of individual blobs *B* associated with the placement of each query on the scaffold tree. For example, the placement in Figure 1B will result in *E* containing edges {2, 5} and {5, 6} and *B* being empty. Note that in many cases, the underlying placement algorithm will suggest multiple alternative placement edges, each with an associated probability for each query. The user has the option to limit the number of placements per query to the top *k* edges with the highest probabilities.

##### Blob placement

For each blob-edge in *E* and each blob in *B*, Splicer identifies the set of query sequences that need to be placed within that subtree and carries out the placement.

##### Placement aggregation

If a query *q* was placed on scaffold edge *e* with probability *p*_1_, and then was placed on a reference tree edge within the corresponding subtree with probability *p*_2_, then the overall placement probability on that edge is *p*_1_ · *p*_2_.

Since some edges are repeated across different blobs and blob-edges, it may be possible that placement of the same query on different scaffold-edges leads to the final placement on the same reference tree edge. In this case, the individual placement probabilities from different placement paths are summed up to produce the final probability for that edge. The aggregated placement probabilities are saved to the *splicer*.*jplace* file.

#### 2.1.3 Proof of Splicer’s accuracy

We show that Splicer can scale any placement algorithm to large trees while guaranteeing that a correct placement edge can always be found.

##### Theorem 2.

*Let P be an accurate placement algorithm (i*.*e*., *it always finds a correct placement edge); then Splicer, when using P as a subroutine, is guaranteed to be accurate*.

*Proof*. We begin by introducing the necessary notation. Let *q* denote the query sequence and let *T* ^∗^ denote the true tree topology over the taxon set *L* ∪ {*q*}, where *L* is the set of reference taxa as before. Let *T* be the reference tree (i.e., the restriction of *T* ^∗^ to *L*) and *S* ⊂ *L* be the scaffold taxa determined by Splicer after decomposing *T* . For a node *x* of *T*, we denote the blob to which *x* belongs by *b*(*x*).

Consider the tree *T* ^∗^|_*S*∪{*q*}_ that is a restriction of *T* ^∗^ to the scaffold taxa and the query sequence. Let (*u, v*) denote the true placement edge of *q* onto the scaffold tree *T* |_*S*_. That is, the parent of *q* subdivides (*u, v*) in *T* ^∗^|_*S*∪{*q*}_. If (*u, v*) is also an edge in *T*, then we are done (because in this case *b*(*u*) and *b*(*v*) are empty and the placement algorithm stops). Assume now that it is not the case and let (*w, x*) be the true placement edge for *q* in *T* . Nodes *w* and *x* must either be themselves on the path from *u* to *v* in *T* or their ancestor is on that path (Figure 2). Note that since none of the intermediate nodes on the path from *u* to *v* in *T* are maintained in *T* |_*S*_, those nodes and their children ‘off’ the *u* − *v* path mush have been absorbed into the *b*(*u*) or *b*(*v*) blobs. Therefore, edge (*w, v*) will be present in the blob-edge {*b*(*u*), *b*(*v*)} if *b*(*u*) ≠ *b*(*v*) or in the blob *b*(*u*) if *b*(*u*) = *b*(*v*).

**Figure 2.**
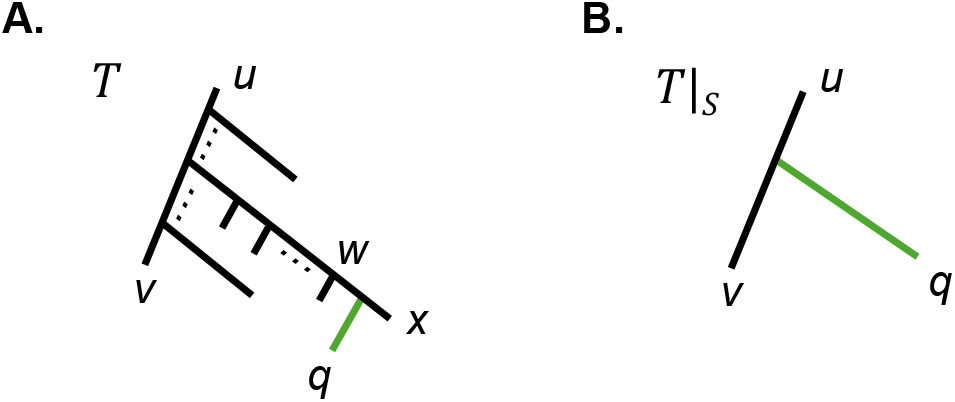
Demonstrating why Splicer does not lose accuracy when performing a two-step placement. **A:** The correct placement of query *q* on the full tree *T* on edge (*w, x*). **B:** Placement on the scaffold tree *T* |_*S*_ with some of the intermediate nodes absorbed into blobs *b*(*u*) and/or *b*(*v*). Splicer will first find the scaffold edge (*u, v*). Then, since the correct edge (*w, x*) is guaranteed to be present within the blob-edge {*b*(*u*), *b*(*v*)}, it will be found in the second placement step.

### 2.2 Automatic sequence classification with Splicer

To make Splicer directly applicable for sequence classification tasks, we implemented a capability to parse a clades definition file during decomposition and automatically assign the most likely clades to new sequences at the placement step. For clade-name formatting, we follow a pattern standard in pathogen surveillance; see, for example, Pango [18], Nextclade [1], and octoFLU [7]. In this format, clades have names of type X.Y.Z (e.g., BA.1.1.7 or 2.3.4.4b), where X is ancestral to X.Y, and X.Y is ancestral to X.Y.Z, and so on. Given a clades definition file for reference sequences, the clade-names are propagated from the leaves to the root of the reference tree according to the following rules (see Figure 3 for an example):

**Figure 3.**
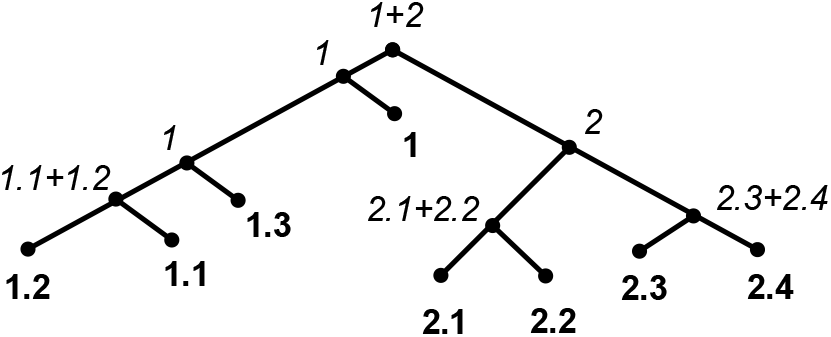
An example of clade-name inference for internal nodes from the clade definitions at the sequence/leaf level. Query placement on a branch leading to a node *v* will get a clade assignment based on the inferred clade-label at *v*.

(i) For an internal node *v*, let 𝒞 be the list of unique clade names below it. We refine 𝒞 so that it contains only minimal clades; i.e., if 𝒞 contains clades ‘X.Y’ and ‘X’, we only keep ‘X’. We denote the refined set by 𝒞^′^
(ii) If |𝒞^′^| = 1, then the clade for *v* is uniquely determined.
(ii) If |𝒞^′^| = 2, then we assign ‘X+Y’ as the clade name for *v* if X and Y are the two clade-names in 𝒞^′^.
(iii) If |𝒞^′^| *>* 2, then we try to find the maximal clade name that is ancestral to all clades in 𝒞^′^. E.g., if 𝒞^′^ contains clades ‘X.Y.U’, ‘X.Y.W’, and ‘X.Y.Z’, then we assign clade ‘X.Y’ to *v*. If no common ancestral clade exists, then we assign ‘*mixed*’ as the clade-name for *v*.

It is not difficult to see that such label propagation can be performed in linear time over the reference tree. Then, at the placement step, if a query *q* was placed on a directed edge (*u, v*), then *q* is assigned the clade-label of *v*. Note that we preserve the original reference tree rooting during the decomposition step and consequent placement steps.

### 2.3 Splicer evaluation on influenza A virus in swine genomic surveillance data

A dataset of n=14,378 publicly available swine H1 influenza A hemagglutinin (HA) gene sequences collected between 1929–2022 was manually annotated by clade-names as described in [3, 2]. There were 49 distinct clades across three H1 lineages. This dataset was representatively subsampled down to 483 sequences, maintaining clades with at most 10 sequences and subsampling larger clades down to 10 sequences using PARNAS v0.1.6 representative sampling [13]. We refer to the original n=14,378 dataset as the *large* dataset and to the subsampled n=483 set as the *small* dataset.

We evaluated the maximum-likelihood placement algorithms pplacer v1.1.alpha17 and EPA-ng v0.3.8 on the small dataset, and we evaluated Splicer, UShER v0.5.4 [20], and SCAMPP+taxtastic v2.0.0 [21] on the large dataset. Both Splicer and SCAMPP were evaluated in combination with pplacer (Splicer+pplacer and SCAMPP+pplacer) and EPA-ng (Splicer+EPA-ng and SCAMPP+EPA-ng). Since the pplacer algorithm requires an input tree with no 0-length branches, we deduplicated the large dataset and removed sequences that coincided with ancestral nodes when running Splicer+pplacer and SCAMPP+pplacer. The deduplicated dataset contained n=9,525 sequences. To evaluate each method, we performed 1,000 iterations of randomly choosing a sequence from the large dataset, removing it from the reference tree (if needed), placing it back, and determining which clade it was placed into using a custom Python script. If a method returned multiple placement options, we evaluated the placement with the maximum probability. Then, the accuracy of a method was determined as the percentage of cases where the inferred clade was correct.

In addition to accuracy, we measured the placement runtime for each method in two configurations: the runtime when a single sequence is being placed (median runtime over 1,000 random replicates) and the runtime when 100 sequences are being placed at the same time (bulk placement). All runtimes were measured on a standard workstation with a 2.6 GHz 6-Core Intel Core i7 processor and 32GB of RAM.

### 2.4 Applying Splicer to SARS-CoV-2 dataset with millions of sequences

We downloaded the Audacity v1.29 SARS-CoV-2 phylogeny from GISAID with 12,659,102 sequences and associated clade metadata [accessed on June 13, 2025] [19, 11]. Additionally, we downloaded a pre-built masked alignment of 16,498,361 SARS-CoV-2 genomes from GISAID [built on February 4, 2024]. There was a total of n=12,621,172 tips on the Audacity phylogeny that had an associated clade and sequence available. We trimmed the tree to those sequences and removed an additional 1000 randomly selected sequences for testing. That is, the final reference phylogeny had n=12,620,172 tips.

## 3 Results

### 3.1 Splicer improves clade-identification for H1 influenza A virus

Influenza A viruses (IAV) circulating in swine globally are genetically diverse and represent a significant risk to public health [2]. The H1 subtype of the hemagglutinin gene of swine IAV is particularly diverse, with approximately 50 genetically and antigenically distinct clades that circulate in swine populations worldwide. The existing approach to classifying new H1 sequences into clades, which informs real-time risk assessment of these viruses, relies on pplacer applied to a reference set of 483 representatively selected sequences deployed on the BV-BRC H1 sequence classifier [17, 3]. To understand the impact of subsampling in sequence classification, we applied Splicer and other scalable placement methods (UShER and SCAMPP) to a full reference dataset of 14,377 sequences, and compared the accuracy of the scalable methods on the full dataset against the accuracy of pplacer and EPA-ng on the small reference dataset.

Figure 4 shows that using the full reference dataset in combination with scalable placement methods significantly improves the classification accuracy of H1 genetic sequences. Splicer showed the highest classification accuracy across all compared methods, with Splicer+EPA-ng and Splicer+pplacer achieving 99.6% and 99.5% accuracy, respectively. The error rate for Splicer+EPA-ng was 10-fold lower than the error rate of EPA-ng on the small reference dataset.

**Figure 4.**
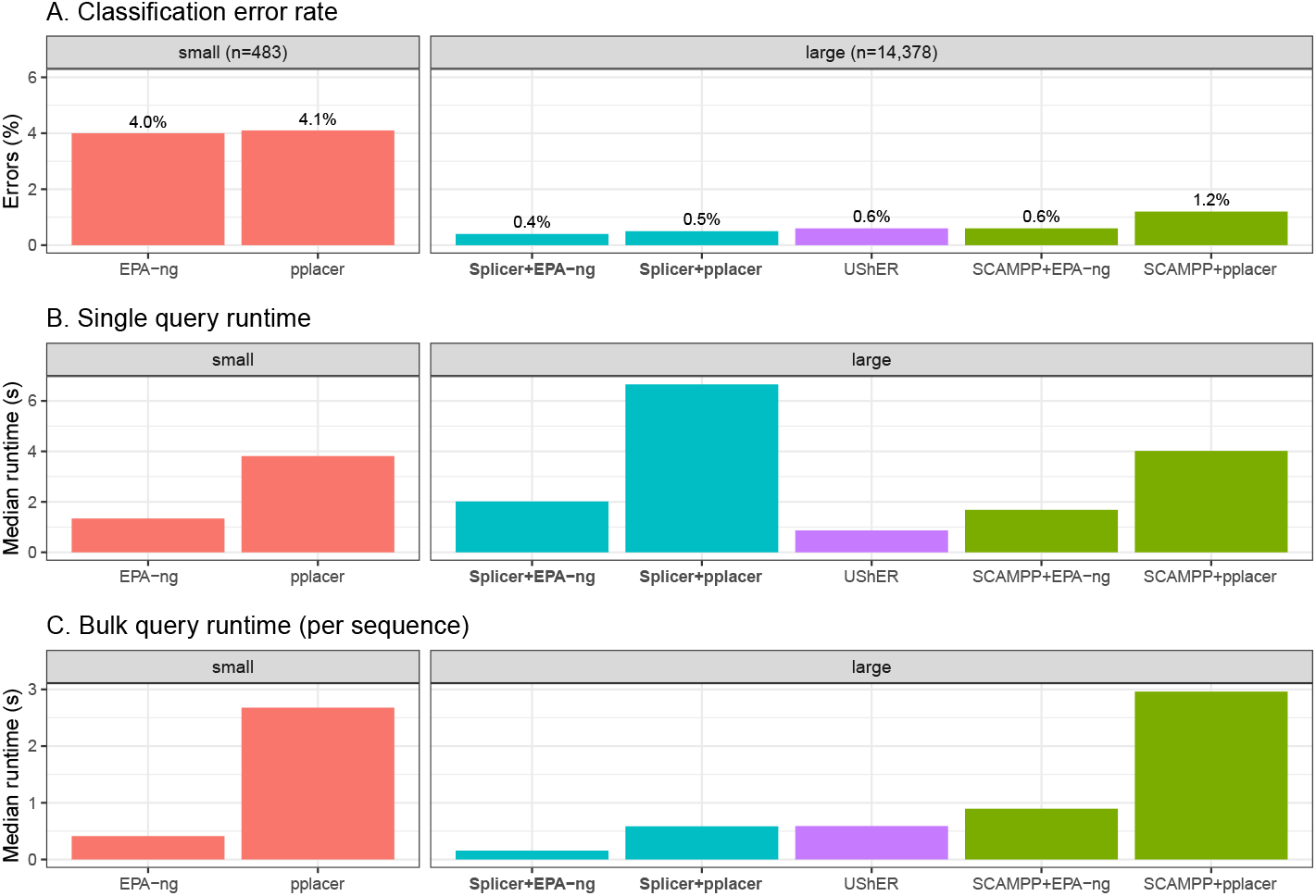
Error rate and runtime of different placement methods on small and large swine influenza A virus H1 reference datasets. **A**: The percentage of test sequences that were incorrectly classified based on their placement. **B**: Median runtime of the placement algorithm when a single query sequence was placed at a time. **C**: Bulk runtime comparison, when 100 query sequences were placed at once.

In terms of the runtime over a single query sequence on the large dataset, UShER was the fastest method with around 0.9 seconds per placement, followed by SCAMPP+EPA-ng and Splicer+EPA-ng, each taking approximately 2 seconds per placement. However, when performing bulk placement of 100 sequences at once, Splicer+EPA-ng was the fastest method, taking only 0.2 seconds per sequence, followed by Splicer+pplacer and UShER at 0.6 seconds per sequence. Notably, Splicer+EPA-ng was significantly faster at bulk placement on the large dataset than base EPA-ng on the small dataset. Similarly, Splicer+pplacer was significantly faster than pplacer at bulk placement, despite differences in dataset size, showing that Splicer is particularly efficient at scaling maximum-likelihood placement for bulk queries.

### 3.2 Splicer scales maximum-likelihood placement to full SARS-CoV-2 dataset

Building and maintaining comprehensive SARS-CoV-2 phylogenies with millions of sequences presented a significant computational challenge throughout the COVID-19 pandemic [16]. This challenge necessitated the development of novel scalable phylogenetic tools, such as UShER [20] and MAPLE [9, 12]. To the best of our knowledge, UShER remains the only placement tool efficiently applicable at a scale of over 10 million SARS-CoV-2 genomes. UShER is used to maintain the only publicly available comprehensive SARS-CoV-2 phylogeny, Audacity, which contains over 12 million genomes. We show that Splicer can, for the first time, scale maximum-likelihood placement to be applicable to the Audacity phylogeny.

We used a single node of the USDA SCINet Ceres computational cluster to run Splicer. The node had 128 available cores and a total of 2,304 GB of RAM. The Splicer decomposition of a reference phylogeny with n=12,620,172 sequences required 4 hours, with the decomposition itself taking 3 hours (1 process, 70GB of RAM) and the remainder of the time spent on I/O operations for saving the decomposition. After the decomposition, the edge lengths within all blobs and the scaffold tree were re-inferred using RAxML-ng v.1.2.2 to enable maximum-likelihood placement, since UShER computes discrete edge lengths via maximum parsimony. One thousand hold-out SARS-CoV-2 genomes were then placed on the decomposed backbone using Splicer+EPA-ng. The placement took a total of 28 hours, i.e., approximately 102 seconds per sequence. Note that we could not run SCAMPP for comparison, as inferring ML branch lengths over the full 12M tree is not feasible, and SCAMPP requires reading the full tree and alignment file for each placement, unlike Splicer.

For the 1,000 hold-out sequences, the clade classifications by Splicer and the clades prescribed by Audacity were identical in 81.7% of cases. Splicer was often more conservative in its clade assignments than UShER, placing sequences in more ancestral clades (13.9% of cases); Splicer chose a more refined clade than UShER/Audacity in 1.2% of cases. Additionally, the two methods prescribed different (incompatible) clades in 3.2% of cases. This comparison suggests that there are likely to be topological differences between maximum likelihood and maximum parsimony SARS-CoV-2 phylogenies if they were built from scratch. Splicer can therefore play a role in building maximum likelihood alternatives to the UShER SARS-CoV-2 phylogenies.

## 4 Discussion

In this work, we introduced the first sub-linear algorithm for phylogenetic placement, Splicer. Splicer can perform placement in 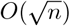 time, where *n* is the number of reference sequences. We proved that if paired with an accurate ‘base’ placement method, Splicer is also guaranteed to be accurate. Based on this observation, we paired Splicer with EPA-ng and pplacer by default, the two most popular and accurate currently available placement methods. In principle, however, Splicer can be adapted to scale-up any placement algorithm, including very fast ones such as UShER. This has significant utility as UShER scales linearly as sequence volume grows and it is currently integrated within existing genomic epidemiology pipelines and public databases.

Using a diverse swine influenza A virus dataset, we showed that Splicer is very accurate and is particularly efficient at bulk placement of multiple query sequences at once (Figure 4). Splicer paired with EPA-ng was a particularly fast combination, taking only 0.2 seconds per query sequence on a reference dataset with over 14,000 references. Additionally, we applied Splicer to a very large SARS-CoV-2 reference dataset comprising over 12 million reference genomes and demonstrated that Splicer can perform maximum-likelihood placement in a reasonable time using the largest publicly available genomic dataset that is used to identify and track viral evolution. This is an important step, as previously, maximum likelihood placement was not feasible at this scale, thereby reducing its potential for implementation in real-time genomic epidemiology.

The current implementation of Splicer is static and is primarily best suited for sequence classification tasks against a predefined taxonomy. The future development of the method will make it dynamic, allowing it to update the reference phylogeny whenever new sequences are placed/added. Conceptually, such an extension is simple, as it requires updating the blob-trees and blob-edges with novel sequences. One can define a threshold maximum blob-size; once this threshold is exceeded for one of the blobs, the decomposition of the reference tree will be recomputed. The concept is similar to the dynamic allocation of space for vectors or hash tables, which is standard in many programming languages, such as Java and Python.

Another possible area of development is further improving the scalability of the method. It is possible to ‘chain’ the decomposition of the reference tree, where, first, a smaller blob size is chosen, and the intermediate scaffold tree is relatively large, and then the intermediate scaffold tree is further decomposed into blobs. This way, one can design nested decompositions with an overall placement runtime of order *n*^1*/*3^, *n*^1*/*4^, and so on. However, such a nested approach represents a challenge in terms of maintaining the accuracy of placement. In particular, if the placement on any of the nested scaffold trees is inaccurate, the following placements will explore the wrong blobs. That is, increasing the number of chained placements increases the probability of placement error. Overcoming this error potential will require exploring neighboring blobs to increase the probability of finding the correct placement edge.

## Code availability

Splicer tool and code are available at https://github.com/flu-crew/splicer.

## Footnotes

1 Although similar in concept, this is not the same as the tree of blobs (TOB) concept used in phylogenetic network literature, where blobs represent bi-connected components of a graph.

## References

[1] Ivan Aksamentov, Cornelius Roemer, Emma B Hodcroft, and Richard A Neher. Nextclade: clade assignment, mutation calling and quality control for viral genomes. Journal of open source software, 6(67):3773, 2021.

[2] Tavis K Anderson, Jennifer Chang, Zebulun W Arendsee, Divya Venkatesh, Carine K Souza, J Brian Kimble, Nicola S Lewis, C Todd Davis, and Amy L Vincent. Swine influenza A viruses and the tangled relationship with humans. Cold Spring Harbor perspectives in medicine, page a038737, 2020.

[3] Tavis K. Anderson, Catherine A. Macken, Nicola S. Lewis, Richard H. Scheuermann, Kristien Van Reeth, Ian H. Brown, Sabrina L. Swenson, Gaëlle Simon, Takehiko Saito, Yohannes Berhane, Janice Ciacci-Zanella, Ariel Pereda, C. Todd Davis, Ruben O. Donis, Richard J. Webby, and Amy L. Vincent. A phylogeny-based global nomenclature system and automated annotation tool for H1 hemagglutinin genes from swine influenza A viruses. mSphere, 1(6), 2016.

[4] Metin Balaban, Shahab Sarmashghi, and Siavash Mirarab. Apples: Fast distance-based phylogenetic placement. In RECOMB, pages 287–288, 2019.

[5] Pierre Barbera, Alexey M Kozlov, Lucas Czech, Benoit Morel, Diego Darriba, Tomáš Flouri, and Alexandros Stamatakis. Epa-ng: massively parallel evolutionary placement of genetic sequences. Systematic biology, 68(2):365–369, 2019.

[6] Simon A Berger, Denis Krompass, and Alexandros Stamatakis. Performance, accuracy, and web server for evolutionary placement of short sequence reads under maximum likelihood. Systematic biology, 60(3):291–302, 2011.

[7] Jennifer Chang, Tavis K Anderson, Michael A Zeller, Phillip C Gauger, and Amy L Vincent. octoFLU: automated classification for the evolutionary origin of influenza A virus gene sequences detected in US Swine. Microbiology resource announcements, 8(32), 2019.

[8] Lucas Czech, Alexandros Stamatakis, Micah Dunthorn, and Pierre Barbera. Metagenomic analysis using phylogenetic placement—a review of the first decade. Frontiers in Bioinformatics, 2:871393, 2022.

[9] Nicola De Maio, Prabhav Kalaghatgi, Yatish Turakhia, Russell Corbett-Detig, Bui Quang Minh, and Nick Goldman. Maximum likelihood pandemic-scale phylogenetics. Nature Genetics, 55(5):746–752, 2023.

[10] Elizabeth Koning, Malachi Phillips, and Tandy Warnow. pplacerDC: a new scalable phylogenetic placement method. In Proceedings of the 12th ACM International Conference on Bioinformatics, Computational Biology, and Health Informatics, pages 1–9, 2021.

[11] R. Lanfear. A global phylogeny of hCoV-19 sequences from GISAID, 2020.

[12] Nhan Ly-Trong, Chris Bielow, Nicola De Maio, and Bui Quang Minh. Cmaple: efficient phylogenetic inference in the pandemic era. Molecular Biology and Evolution, 41(7):msae134, 2024.

[13] Alexey Markin, Sanket Wagle, Siddhant Grover, Amy L Vincent Baker, Oliver Eulenstein, and Tavis K Anderson. Parnas: objectively selecting the most representative taxa on a phylogeny. Systematic Biology, 72(5):1052–1063, 2023.

[14] Frederick A Matsen, Robin B Kodner, and E Armbrust. pplacer: linear time maximumlikelihood and bayesian phylogenetic placement of sequences onto a fixed reference tree. BMC bioinformatics, 11(1):1–16, 2010.

[15] Adam Monier, Jean-Michel Claverie, and Hiroyuki Ogata. Taxonomic distribution of large dna viruses in the sea. Genome biology, 9(7):R106, 2008.

[16] Benoit Morel, Pierre Barbera, Lucas Czech, Ben Bettisworth, Lukas Hübner, Sarah Lutteropp, Dora Serdari, Evangelia-Georgia Kostaki, Ioannis Mamais, Alexey M Kozlov, Pavlos Pavlidis, Dimitrios Paraskevis, and Alexandros Stamatakis. Phylogenetic analysis of SARS-CoV-2 data is difficult. Molecular Biology and Evolution, 38(5):1777–1791, 12 2020.

[17] Robert D Olson, Rida Assaf, Thomas Brettin, Neal Conrad, Clark Cucinell, James J Davis, Donald M Dempsey, Allan Dickerman, Emily M Dietrich, Ronald W Kenyon, et al. Introducing the bacterial and viral bioinformatics resource center (bv-brc): a resource combining patric, ird and vipr. Nucleic acids research, 51(D1):D678–D689, 2023.

[18] Andrew Rambaut, Edward C Holmes, Áine O’Toole, Verity Hill, John T McCrone, Christopher Ruis, Louis du Plessis, and Oliver G Pybus. A dynamic nomenclature proposal for sars-cov-2 lineages to assist genomic epidemiology. Nature microbiology, 5(11):1403–1407, 2020.

[19] Yuelong Shu and John McCauley. GISAID: Global initiative on sharing all influenza data–from vision to reality. Eurosurveillance, 22(13):30494, 2017.

[20] Yatish Turakhia, Bryan Thornlow, Angie S Hinrichs, Nicola De Maio, Landen Gozashti, Robert Lanfear, David Haussler, and Russell Corbett-Detig. Ultrafast sample placement on existing trees (usher) enables real-time phylogenetics for the sars-cov-2 pandemic. Nature Genetics, 53(6):809–816, 2021.

[21] Eleanor Wedell, Yirong Cai, and Tandy Warnow. Scampp: scaling alignment-based phylogenetic placement to large trees. IEEE/ACM Transactions on Computational Biology and Bioinformatics, 20(2):1417–1430, 2022.

